# Green Synthesized Zinc Oxide Nanoparticles from *Azadirachta indica* Exhibit Enhanced Antibacterial, Antioxidant and Cytotoxic Activities

**DOI:** 10.64898/2026.08.12.744370

**Authors:** Sadia Manzoor, Tayyaba Arif, Hamza Rafiq, Saima Younas, Shahina Akter

## Abstract

Green synthesis of zinc oxide nanoparticles (ZnO NPs) offers a sustainable strategy for developing multifunctional antimicrobial nanomaterials. In this study, ZnO NPs were synthesized using *Azadirachta indica* leaf extract and characterized by UV-vis spectroscopy, FTIR, XRD, SEM, and GC-MS. The nanoparticles exhibited a characteristic absorption peak at 352 nm, a direct band gap of 3.07 eV, and hexagonal wurtzite crystallinity with an average crystallite size of approximately 32 nm. The biosynthesized ZnO NPs showed concentration-dependent antibacterial activity against *Erwinia carotovora*, producing inhibition zones of up to 25.9 mm. Mechanistic studies revealed significant membrane damage, evidenced by 4.77-fold and 5.62-fold increases in extracellular protein and amino acid leakage, respectively, with marked alterations in bacterial protein profiles detected by SDS-PAGE. The nanoparticles also exhibited strong antioxidant activity, achieving 89.4% DPPH radical scavenging, and induced dose-dependent cytotoxicity in HepG2 cells with an estimated IC_50_ of 124.8 μg/mL. These findings demonstrate that neem-mediated ZnO nanoparticles possess potent antibacterial activity through membrane disruption while exhibiting promising antioxidant properties, highlighting their potential as eco-friendly nanomaterials for the management of bacterial soft rot and other phytopathogenic diseases.

## 1. Introduction

Nanotechnology has transformed the development of functional materials by enabling precise control over physicochemical properties at the nanoscale. Compared with the classical bulk application, nanomaterials possess a larger surface-to-volume ratio, unique optical and electronic characteristics, and enhanced surface reactivity, making them attractive for applications ranging from catalysis and environmental remediation to biomedicine and agriculture (Dey et al., 2025). Among metal oxide nanomaterials, zinc oxide (ZnO) nanoparticles have attracted particular interest because of their wide direct band gap, high exciton binding energy, chemical stability and broad-spectrum antimicrobial activity (Lebaka et al., 2025; Takcı et al., 2025). Zinc is also an essential micronutrient for plants, while zinc oxide has received Generally Recognized As Safe (GRAS) status for specific applications, supporting its continued investigation for biomedical and agricultural uses (Rani et al., 2025).

The physicochemical properties of ZnO nanoparticles are strongly influenced by their synthesis strategy. Conventional physical and chemical methods provide good control over particle formation but frequently rely on toxic reagents, hazardous solvents and energy-intensive processing, raising concerns regarding environmental sustainability and large-scale application (Swain et al., 2025). Green synthesis using plant extracts offers an alternative approach in which naturally occurring phytochemicals act simultaneously as reducing, stabilizing and capping agents during nanoparticle formation (Al-darwesh et al., 2024; Villagrán et al., 2024). Besides reducing chemical waste, these phytochemicals remain associated with the nanoparticle surface and can modify surface chemistry, colloidal stability and biological performance (El-Saadony et al., 2024). The synthesis route becomes an important determinant of the biological behaviour of ZnO nanoparticles.

Among medicinal plants, *Azadirachta indica* (neem) is particularly suitable for green nanoparticle synthesis because its leaves are rich in flavonoids, polyphenols, terpenoids, azadirachtin, nimbolide and salannin, compounds known for their antimicrobial and antioxidant properties (Al-darwesh et al., 2024; Sarkar et al., 2021). These metabolites participate in nanoparticle nucleation and crystal formation while forming a phytochemical coating around the ZnO core, which may further enhance biological activity (Halder et al., 2025; Yagoub et al., 2022). Previous studies have shown that neem-mediated ZnO nanoparticles exhibit well-defined hexagonal wurtzite crystal structures together with improved antimicrobial performance compared with many chemically synthesized counterparts. Such observations suggest that both the intrinsic properties of ZnO and the phytochemical surface layer contribute to biological activity.

Rather than acting through a single antibacterial target, ZnO nanoparticles exert antimicrobial effects through multiple interconnected mechanisms. Initial electrostatic interactions with the bacterial cell envelope compromise membrane integrity and increase permeability, resulting in leakage of intracellular constituents. Simultaneously, the generation of reactive oxygen species (ROS) induces oxidative stress, damaging membrane lipids, proteins, and nucleic acids, while released Zn²⁺ ions further disrupt essential metabolic pathways and enzymatic functions (Mohammed et al., 2025; Nan et al., 2024; Sirelkhatim et al., 2015). The relative contribution of these mechanisms is influenced by particle size, crystallinity and surface chemistry, emphasizing the importance of establishing relationships between structure and activity when evaluating newly synthesized nanomaterials.

*Erwinia carotovora* (*Pectobacterium carotovorum*), the causal agent of bacterial soft rot, causes extensive postharvest and field losses in potato, tomato and several horticultural crops through secretion of cell wall-degrading enzymes that rapidly macerate plant tissues (Fei et al., 2026; Perfileva et al., 2025). The pathogen survives in plant debris, in the soil and water and is spread by contaminated material and insect vectors making management difficult. Despite advances in green nanoparticle synthesis, most published studies have emphasized nanoparticle preparation and conventional antibacterial assays, whereas fewer studies are performed on the direct application of these chemicals to phytopathogenic bacteria and check their membrane disruption, intracellular leakage, protein profile alterations and phytochemical composition. The present study investigated neem-mediated ZnO nanoparticles synthesized through a green approach and examined how their physicochemical characteristics relate to antibacterial performance against *E. carotovora*. Structural and chemical characterization was performed using UV-Vis spectroscopy, FTIR, X-ray diffraction, SEM, TEM and GC-MS analyses. Antibacterial activity was evaluated through complementary biochemical and microbiological assays, including DPPH radical scavenging, Bradford protein leakage, ninhydrin amino acid leakage, SDS-PAGE protein profiling, zone of inhibition and MTT cytotoxicity. Integrating these complementary datasets provides a broader understanding of the structure-activity relationship underlying the antibacterial behaviour of neem-mediated ZnO nanoparticles and supports their development as sustainable antimicrobial materials for agricultural applications.

## 2. Materials and Methods

### 2.1 Materials

Fresh leaves of *A. indica* were collected from the Centre for Applied Molecular Biology, University of Lahore, Lahore, Pakistan, in March 2025 and authenticated by the Department of Botany (Voucher No. ZnONEEM2025001). Zinc sulfate heptahydrate (ZnSO₄·7H₂O, 99% purity), sodium hydroxide (NaOH), methanol, ethanol, gentamicin sulfate, 2,2-diphenyl-1-picrylhydrazyl (DPPH), ninhydrin, bovine serum albumin (BSA), Dulbecco’s Modified Eagle Medium (DMEM), fetal bovine serum (FBS), penicillin, streptomycin and other analytical-grade reagents were obtained from Sigma-Aldrich (St. Louis, MO, USA). Deionized water was used throughout the study.

### 2.2 Preparation of neem leaf extract

Fresh neem leaves were thoroughly washed with distilled water, shade-dried under controlled conditions (25°C; relative humidity 40–60%) for 14 days and then homogenized using mortar and pestle. An aqueous extract was prepared by boiling 10 g of leaf powder in 100 mL of deionized water at 60°C for 10 min with continuous stirring (Tsegahun & Aklilu, 2025). After cooling, the extract was filtered through Whatman No. 1 filter paper, and the filtrate was used immediately for nanoparticle synthesis.

### 2.3 Green synthesis of ZnO nanoparticles

Neem-mediated ZnO nanoparticles were synthesized using a modified green precipitation method based on reported protocols (Pachaiappan et al., 2021). Briefly, 35 mL of 0.1 M ZnSO₄·7H₂O solution was stirred continuously, followed by the dropwise addition of 20 mL of freshly prepared neem extract. The pH was adjusted to approximately 12 using 2 M NaOH, and the reaction mixture was maintained at 80°C for 30 min under continuous stirring until a yellow-brown suspension was obtained, indicating nanoparticle formation. The precipitate was collected by centrifugation (10,000 × g, 20 min), washed three times with deionized water followed by absolute ethanol, dried at 60°C, finely powdered, and stored in amber glass vials at 4°C until further analysis.

### 2.4 Physicochemical characterization of ZnO nanoparticles

The optical properties of the synthesized ZnO nanoparticles were characterized using a Cary 60 UV–Visible spectrophotometer (Agilent Technologies, Santa Clara, CA, USA) over the wavelength range of 200–600 nm. The optical band gap was estimated from the Tauc plot assuming a direct electronic transition. Surface functional groups involved in nanoparticle formation and phytochemical capping were identified by Fourier-transform infrared (FTIR) spectroscopy over the spectral range of 400–4000 cm⁻¹. Crystal structure and phase purity were analysed using a Rigaku MiniFlex 600 X-ray diffractometer equipped with Cu Kα radiation (λ = 0.154 nm) (Rahman et al., 2022). Crystallite size was estimated using the Debye–Scherrer equation. The surface morphology of the biosynthesized ZnO nanoparticles was examined using a Nova Nano SEM (FEI, Hillsboro, OR, USA) equipped with a STEM detector and operated at an accelerating voltage of 15 kV. Images were acquired at 200,000× magnification to evaluate particle morphology and aggregation. Particle size distribution was determined by measuring 100 randomly selected, distinguishable nanoparticles from representative SEM micrographs using ImageJ software. Phytochemicals associated with the synthesized nanoparticles were identified by GC-MS using an Agilent 7890B gas chromatograph coupled to an Agilent 5977C mass selective detector fitted with an HP-5 capillary column. Helium was used as the carrier gas (1.0 mL/min), with a 1 μL injection volume and an injector temperature of 250°C. Mass spectra were recorded over an m/z range of 35-498, and compounds were identified using the NIST17 spectral library (Rahman et al., 2022).

### 2.5 Biological evaluation of neem-mediated ZnO nanoparticles

#### 2.5.1 Antibacterial activity

*E. carotovora* (ATCC 39581) was obtained from the University of the Punjab, Lahore, Pakistan, and cultured in Luria–Bertani (LB) broth at 37°C with shaking (200 rpm) for 14 h. The bacterial suspension was adjusted to approximately 1 × 10⁸ CFU/mL (OD₆₀₀ = 0.5). ZnO nanoparticles were dispersed in sterile phosphate-buffered saline (PBS; pH 7.4) by ultrasonication for 10 min and evaluated at concentrations of 50, 100, 150 and 200 μg/mL. The concentration range was selected based on preliminary optimization experiments to evaluate the dose-dependent antibacterial activity. Untreated cultures and gentamicin sulfate (50 μg/mL) served as the negative and positive controls, respectively. Antibacterial activity was determined by the agar disk diffusion method using sterile 6 mm paper disks, and inhibition zone diameters were measured after 24 h of incubation at 37°C (Sharma et al., 2025).

#### 2.5.2 Membrane integrity assays

Membrane damage induced by ZnO nanoparticles was assessed by determining extracellular protein and amino acid leakage. For the Bradford assay, treated bacterial cultures were centrifuged (10,000 × g, 5 min, 4°C), and the supernatant was mixed with Bradford reagent (1:4, v/v). After incubation for 10 min at 25°C, absorbance was measured at 595 nm, and protein concentrations were calculated using a BSA standard curve (Bradford, 1976). Amino acid leakage was quantified using the ninhydrin assay according to Khalil & Villota (1988). Briefly, the cell-free supernatant was reacted with freshly prepared ninhydrin reagent, heated at 95°C for 10 min, diluted with 60% ethanol after cooling, and absorbance was recorded at 570 nm using an L-leucine standard curve for quantification.

#### 2.5.3 SDS–PAGE analysis

Alterations in bacterial protein profiles following nanoparticle exposure were analysed by SDS– PAGE. Bacterial cells were harvested, washed with ice-cold PBS, lysed by sonication in Tris–HCl buffer containing Triton X-100 and SDS, and protein concentrations were normalized using the Bradford assay (Bradford, 1976). Equal amounts of protein (5 μg) were separated on 12% polyacrylamide gels at 120 V. Protein bands were visualized with Coomassie Brilliant Blue R-250, documented using a Bio-Rad Gel Doc XR+ imaging system, and analysed with ImageJ.

#### 2.5.4 Antioxidant activity

The antioxidant activity of neem-mediated ZnO nanoparticles was evaluated using the DPPH radical scavenging assay following Faisal et al. (2021). Nanoparticle suspensions were incubated with DPPH solution for 30 min in the dark, and absorbance was measured at 517 nm (El-beltagi et al., 2024). Radical scavenging activity was calculated according to the following equation;

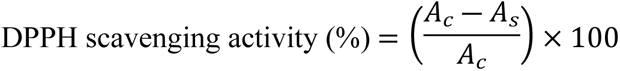

where **Ac** is the absorbance of the control (DPPH solution without nanoparticles) and **As** is the absorbance of the sample containing ZnO nanoparticles. Higher scavenging percentages indicate greater antioxidant activity.

#### 2.5.5 MTT cytotoxicity assay

The cytocompatibility of ZnO nanoparticles was evaluated using HepG2 cells cultured in DMEM supplemented with 10% fetal bovine serum, penicillin (100 IU/mL) and streptomycin (100 μg/mL) at 37°C in a humidified atmosphere containing 5% CO₂. Cells (1 × 10⁴ cells/well) were seeded into 96-well plates, exposed to the respective nanoparticle concentrations for 48 h, and cell viability was determined using the MTT assay as originally described by Mosmann (1983) and following standard procedures for ZnO nanoparticle cytotoxicity assessment in HepG2 cells (Wahab et al., 2014). Absorbance was measured using a Synergy H1 Hybrid microplate reader, and cell viability was calculated using the following equation;

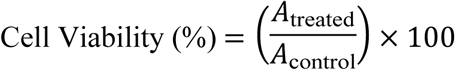

where *A*_treated_ and *A*_control_ represent the mean absorbance values of nanoparticle-treated and untreated control cells, respectively.

### 2.6 Statistical analysis

All experiments were performed using three independent biological replicates, and data are presented as the mean ± standard deviation (SD). Statistical analyses were conducted using GraphPad Prism version 10.0 (GraphPad Software, San Diego, CA, USA). Differences among treatments were analyzed by one-way analysis of variance (ANOVA) followed by Tukey’s multiple comparison test, with statistical significance accepted at P < 0.05.

## 3. Results

### 3.1 Physicochemical characterization of neem-mediated ZnO nanoparticles

#### 3.1.1 UV–Visible spectroscopy and optical band gap

The UV–Visible absorption spectrum of the synthesized ZnO nanoparticles displayed a distinct absorption maximum at 352 nm (Fig. 1A), confirming the formation of ZnO nanoparticles through the green synthesis process. No additional absorption bands were observed within the scanned wavelength range, indicating the absence of detectable secondary products. The optical band gap estimated from the Tauc plot was 3.07 eV (Fig. 1B), consistent with the semiconducting behavior of nanoscale ZnO. The relatively narrow band gap compared with bulk ZnO is indicative of nanoscale crystallites and surface modification by plant-derived constituents.

**Fig. 1.**
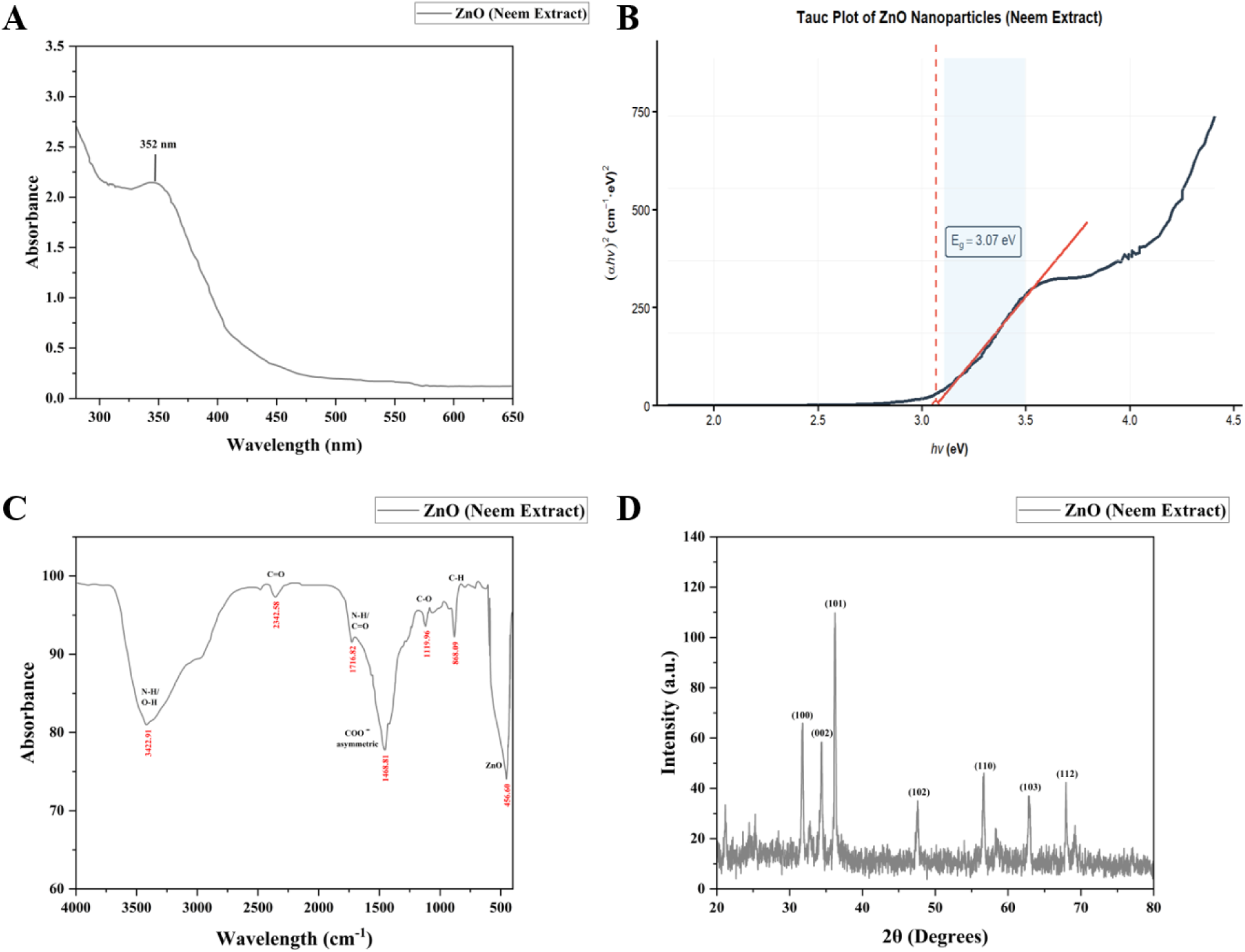
**(A)** UV-visible absorption spectrum, **(B)** Tauc plot of neem-mediated ZnO nanoparticles showing an absorption maximum at approximately 352 nm and an optical band gap of about 3.07 eV, **(C)** FTIR spectrum of neem-mediated ZnO nanoparticles showing phytochemical functional groups and the characteristic Zn-O vibration, **(D)** XRD pattern of neem-mediated ZnO nanoparticles confirming the hexagonal wurtzite ZnO phase and crystalline structure.

#### 3.1.2 FTIR analysis

FTIR spectra revealed several characteristic absorption bands associated with functional groups present on the nanoparticle surface (Fig. 1C). A broad band centered at 3422.91 cm⁻¹ corresponded to O–H/N–H stretching vibrations, while the absorption peak at 1716.82 cm⁻¹ was assigned to C=O stretching. Peaks at 1468.81, 1119.96, and 868.09 cm⁻¹ were attributed to aromatic C–H bending, C–O stretching and C–H vibrations, respectively. A characteristic absorption band at 456.60 cm⁻¹ corresponded to ZnO stretching vibrations, confirming the formation of ZnO nanoparticles. The observed spectrum also indicated the presence of plant-derived functional groups associated with the nanoparticle surface.

#### 3.1.3 X-ray diffraction analysis

The XRD pattern of the synthesized nanoparticles exhibited well-defined diffraction peaks at 31.70°, 34.40°, 36.24°, 47.50°, 56.60°, 62.90° and 67.90°, corresponding to the (100), (002), (101), (102), (110), (103) and (112) crystallographic planes of hexagonal wurtzite ZnO (Fig. 1D). No additional diffraction peaks associated with secondary crystalline phases were detected.

Among the recorded reflections, the (101) plane at 36.24° showed the highest diffraction intensity. Crystallite sizes calculated using the Debye–Scherrer equation ranged from 23.9 to 34.1 nm, with an average crystallite size of approximately 32 nm, confirming the nanoscale crystalline nature of the synthesized ZnO nanoparticles.

#### 3.1.4 SEM Analysis

SEM revealed that the biosynthesized ZnO nanoparticles were predominantly spherical to quasi-spherical and formed moderately agglomerated assemblies (Fig. 2). Individual nanoparticles remained distinguishable within the aggregates, indicating successful nanoparticle formation while reflecting the tendency of nanosized ZnO particles to cluster because of their high surface energy. The particle size distribution determined from SEM micrographs (Supplementary Fig. S1) showed diameters ranging from 34 to 97 nm, with most particles distributed between 50 and 70 nm. The mean particle diameter was 58.9 ± 14.2 nm (n = 100). The relatively uniform morphology and narrow particle size distribution are consistent with controlled particle development during the green synthesis process.

**Fig. 2.**
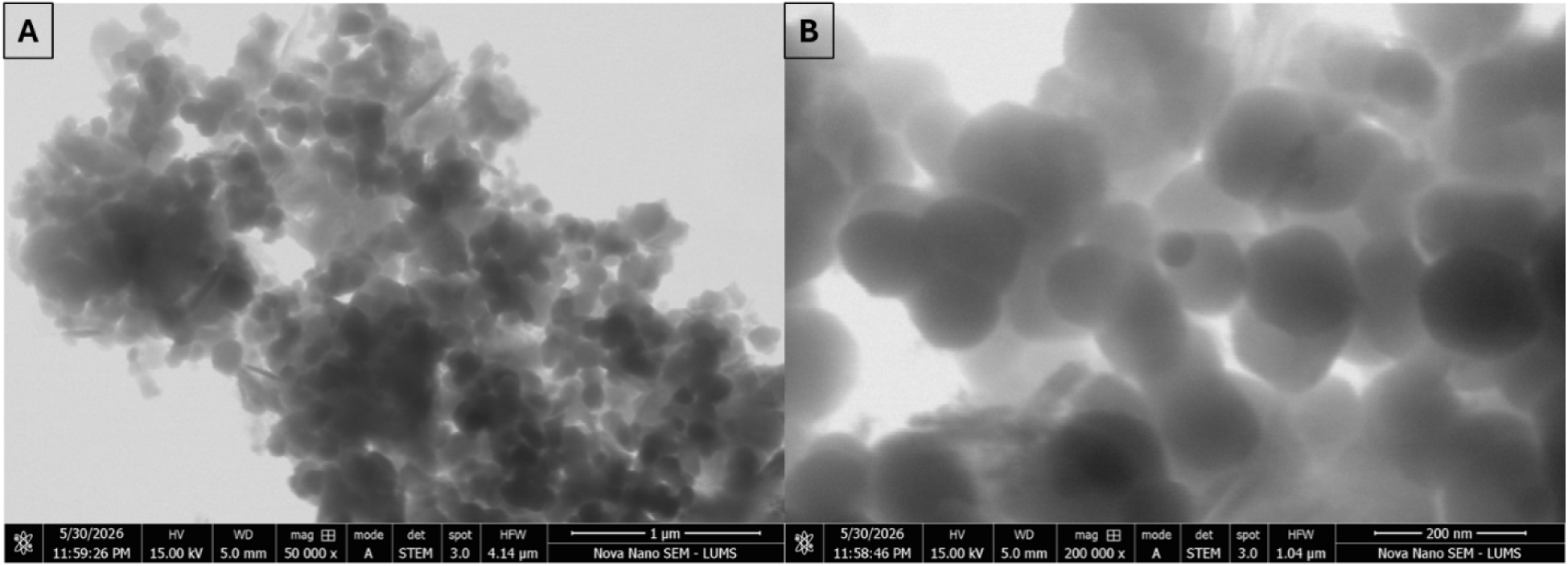
**(A)** 1μm, **(B)** 200nm SEM micrograph of neem-mediated ZnO nanoparticles obtained after green synthesis, illustrating the characteristic surface architecture of the nanoparticle assemblies.

#### 3.1.5 GC-MS phytochemical profiling

GC-MS analysis of the methanolic extract of neem-mediated ZnO nanoparticles detected 17 chromatographic peaks with retention times ranging from 2.06 to 4.25 min (Fig. 3A). The chromatogram indicated the presence of multiple volatile phytochemical constituents associated with the nanoparticle preparation. The most abundant chromatographic peak was observed at 2.498 min, representing 21.96% of the total peak area (Table 1). Compound classification indicated that hydrazine derivatives constituted the predominant class of detected metabolites, followed by acetic acid esters, carbamic acid derivatives, aromatic compounds and hydroxylamine derivatives (Fig. 3B). Compound identities were assigned based on NIST library matching and should be considered tentative pending further structural confirmation.

**Fig. 3.**
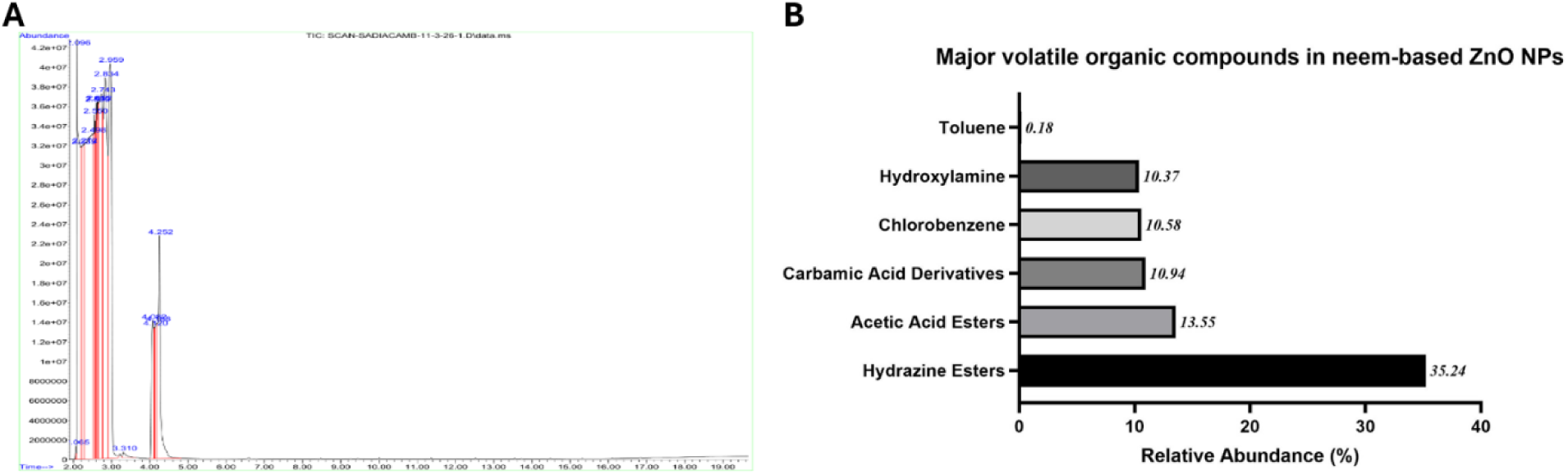
**(A)** GC-MS chromatogram of neem-derived ZnO nanoparticles showing reported peaks between approximately 2.06 and 4.25 min, **(B)** Major volatile organic compounds identified in neem-mediated ZnO nanoparticles by GC–MS analysis, expressed as relative abundance (%).

**Table 1.**
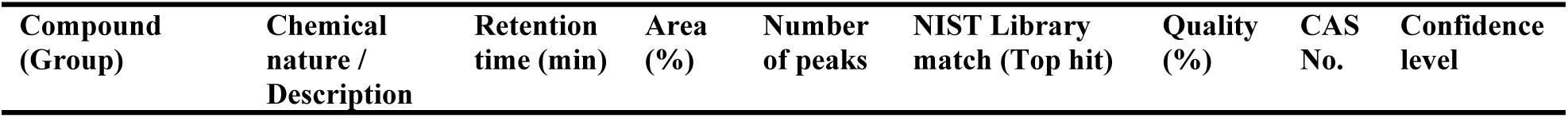
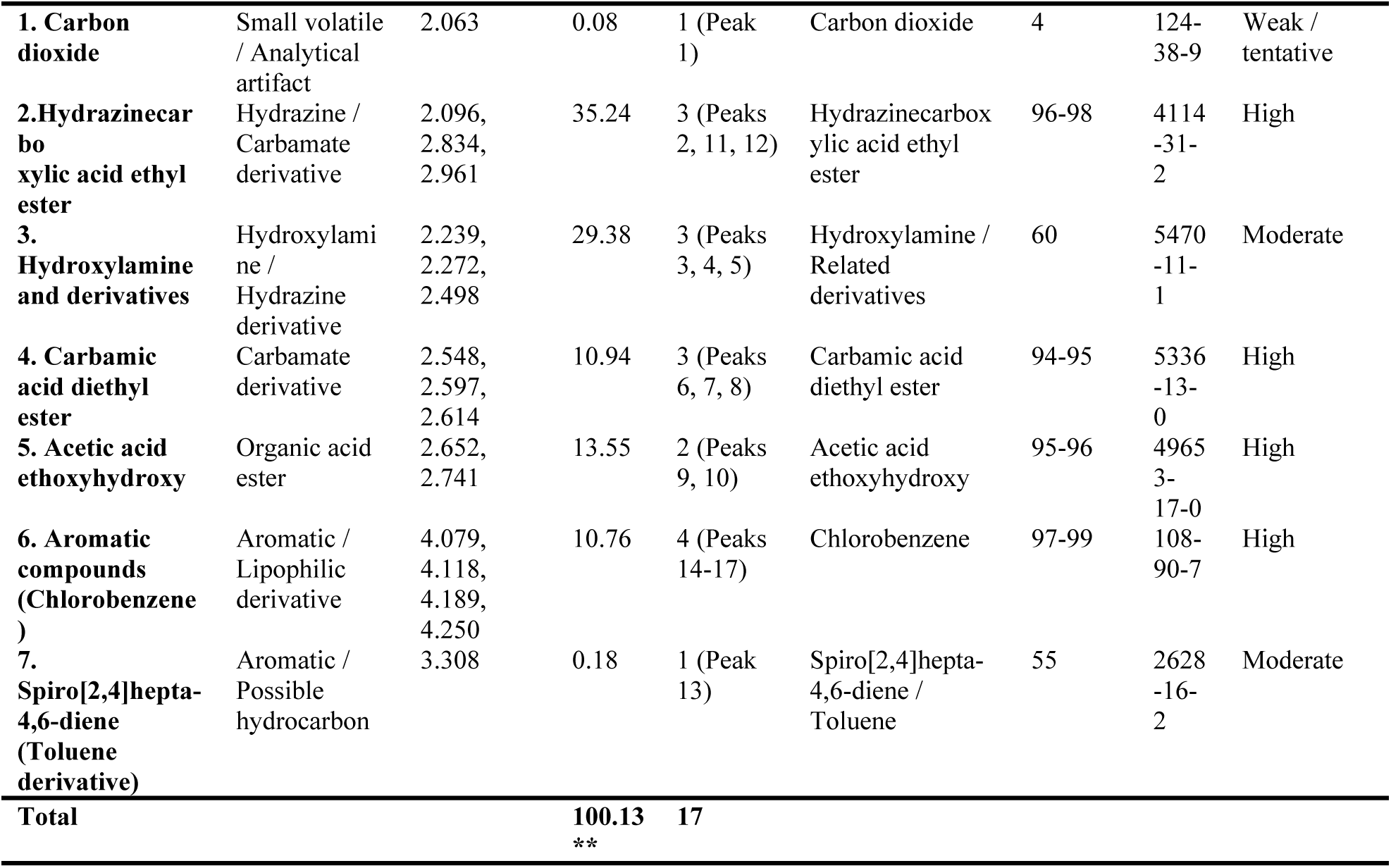
GC-MS-based identification of major volatile compounds in neem-mediated ZnO nanoparticle preparation.

### 3.2 Biological evaluation of neem-mediated ZnO nanoparticles

#### 3.2.1 Antibacterial activity

The antibacterial activity of neem-mediated ZnO nanoparticles against *E. carotovora* was evaluated using the agar disk diffusion assay (Fig. 4A). No inhibition zone was observed in the untreated control, whereas all nanoparticle treatments inhibited bacterial growth in a concentration-dependent manner. The mean inhibition zone increased from 0.92 cm at 50 μg/mL to 1.09 cm, 1.69 cm, and 2.59 cm at 200 μg/mL (Fig. 4B). Gentamicin (50 μg/disk) produced a larger inhibition zone than the nanoparticle treatments and served as the positive control. These results demonstrate a significant concentration-dependent inhibition of *E. carotovora* growth by the synthesized ZnO nanoparticles.

**Fig. 4.**
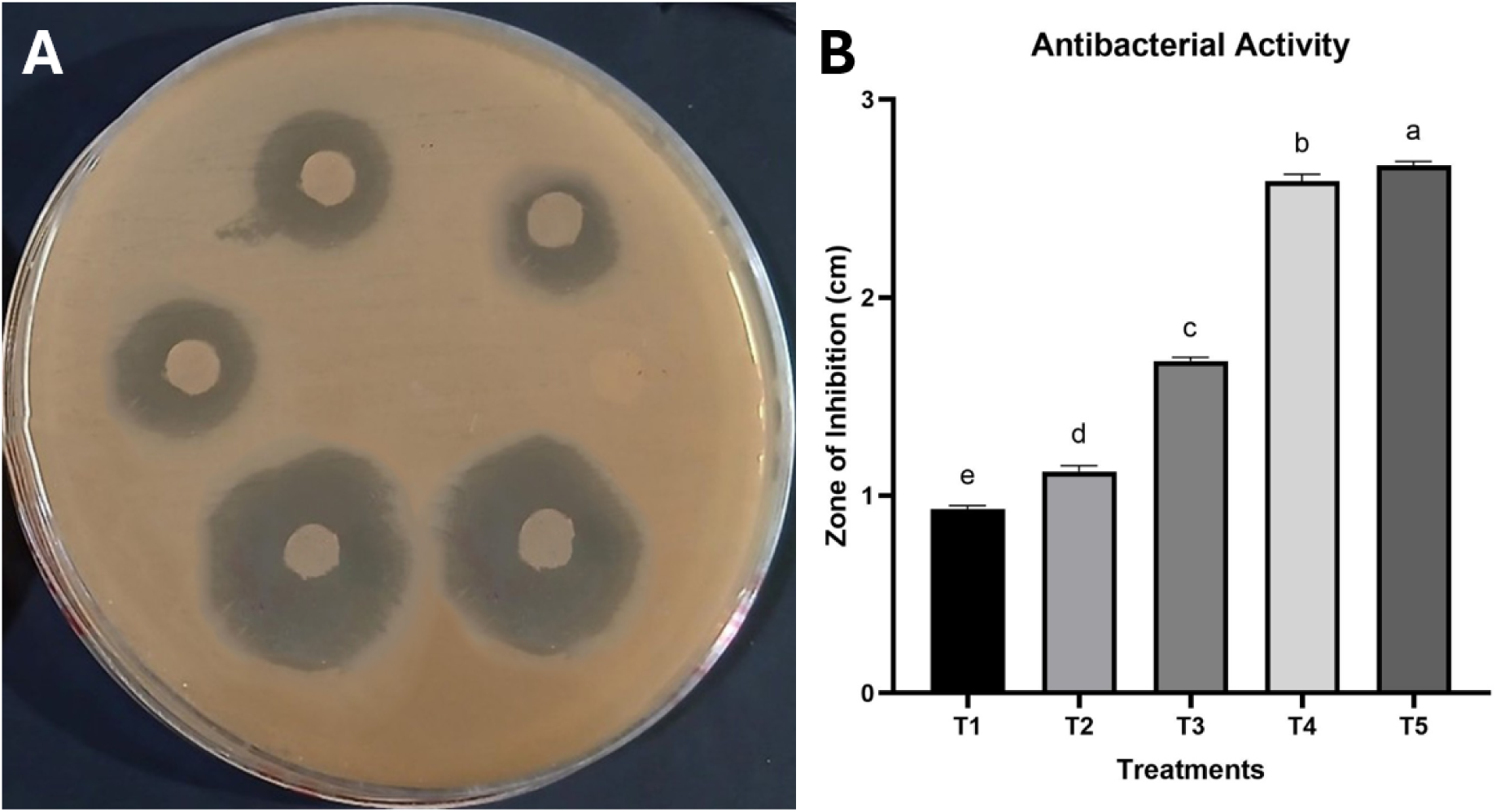
**(A)** Comparative zone of inhibition plates showing concentration-dependent antibacterial activity and control response, **(B)** Antibacterial activity of neem-mediated ZnO nanoparticles against *E. carotovora* determined by the agar well diffusion assay. T1-T4 correspond to ZnO nanoparticle concentrations of 50, 100, 150, and 200 μg/mL, respectively, and T5 represents the positive control (gentamicin, 50 μg/well). Bars represent the mean inhibition zone (cm) ± SD (n = 3). Different lowercase letters indicate statistically significant differences among treatments (P < 0.05, one-way ANOVA followed by Tukey’s HSD test).

#### 3.2.2 Membrane integrity

Exposure of *E. carotovora* to neem-mediated ZnO nanoparticles resulted in a progressive increase in extracellular protein leakage (Fig. 5A). The absorbance at 595 nm increased from 0.131 in the untreated control to 0.220, 0.342, 0.461, and 0.623 following treatment with ZnO nanoparticles at increasing concentrations, representing a 4.77-fold increase relative to the untreated control. The positive control exhibited an absorbance of 0.650. The relationship between nanoparticle concentration and protein leakage was highly linear.

**Fig. 5.**
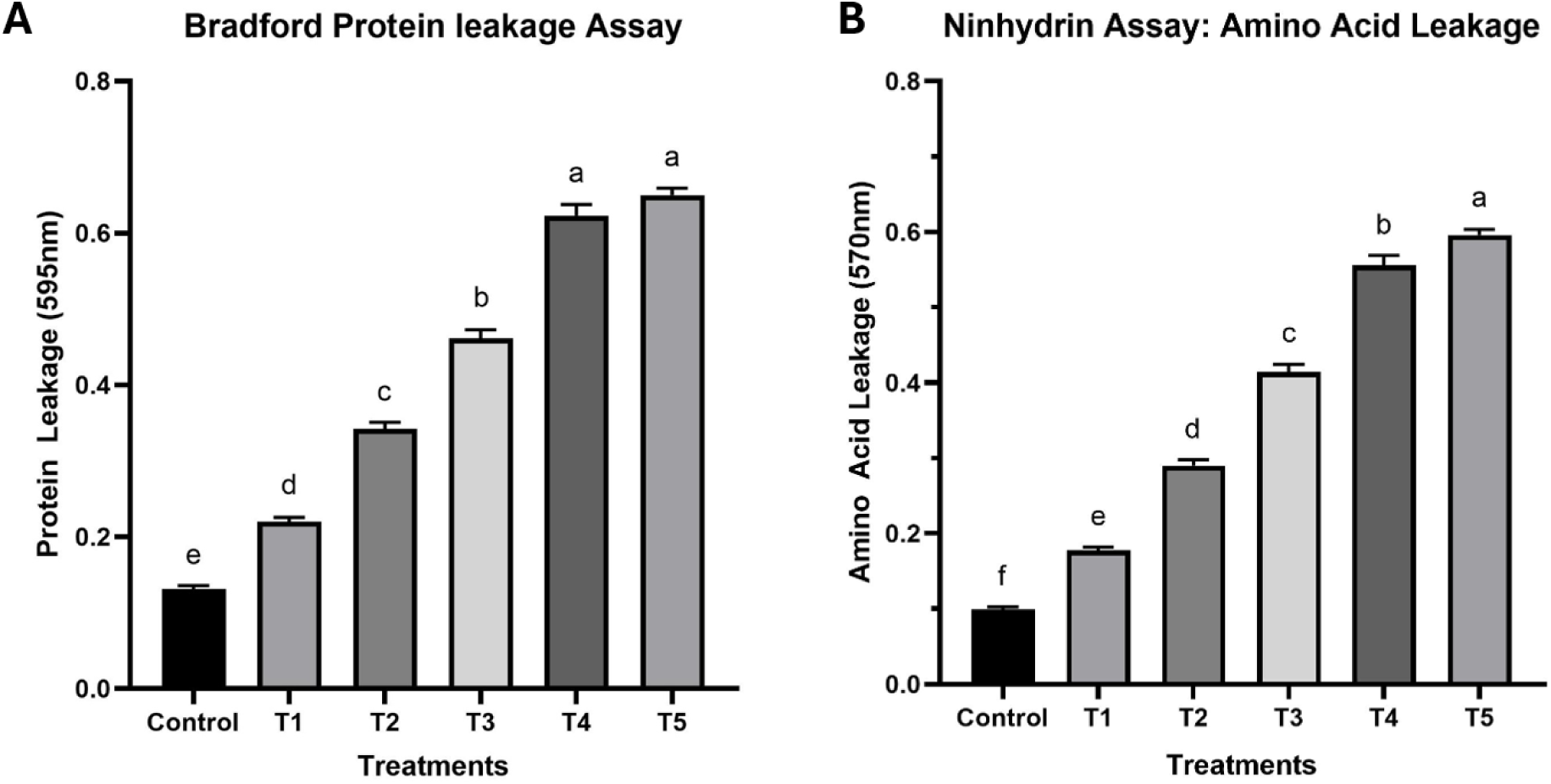
**(A)** Protein leakage, **(B)** Amino acid leakage from *E. carotovora* cells following treatment with neem-mediated ZnO nanoparticles, determined by the Bradford protein assay. The Control represents untreated bacterial cells, while T1-T4 correspond to ZnO nanoparticle concentrations of 50, 100, 150, and 200 μg/mL, respectively, and T5 represents the positive control (gentamicin, 50 μg/well). Bars represent the mean absorbance at 595 nm ± SD (n = 3). Different lowercase letters indicate statistically significant differences among treatments (P < 0.05, one-way ANOVA followed by Tukey’s HSD test).

A similar response was observed for amino acid leakage (Fig. 5B). A concentration-dependent increase in amino acid leakage was observed following treatment with neem-mediated ZnO nanoparticles, with absorbance at 570 nm increasing from 0.099 in the untreated control to 0.177, 0.290, 0.414, and 0.556. The positive control (gentamicin) showed an absorbance of 0.596. At the highest nanoparticle concentration, amino acid leakage was 5.62-fold greater than that of untreated control. The concentration-response relationship was also highly significant.

#### 3.2.3 SDS–PAGE protein profile

SDS–PAGE analysis showed concentration-dependent alterations in the bacterial protein profile following ZnO nanoparticle treatment (Fig. 6). Untreated cells displayed distinct protein bands across the examined molecular weight range. With increasing nanoparticle concentration, band intensity progressively decreased, particularly in the 29 kDa and 66 kDa regions. At the highest treatment level, several bands became faint or were no longer detectable, whereas only minor differences were observed at the lowest treatment concentrations. Diffuse band smearing was also evident within the 20–45 kDa region in highly treated samples.

**Fig. 6.**
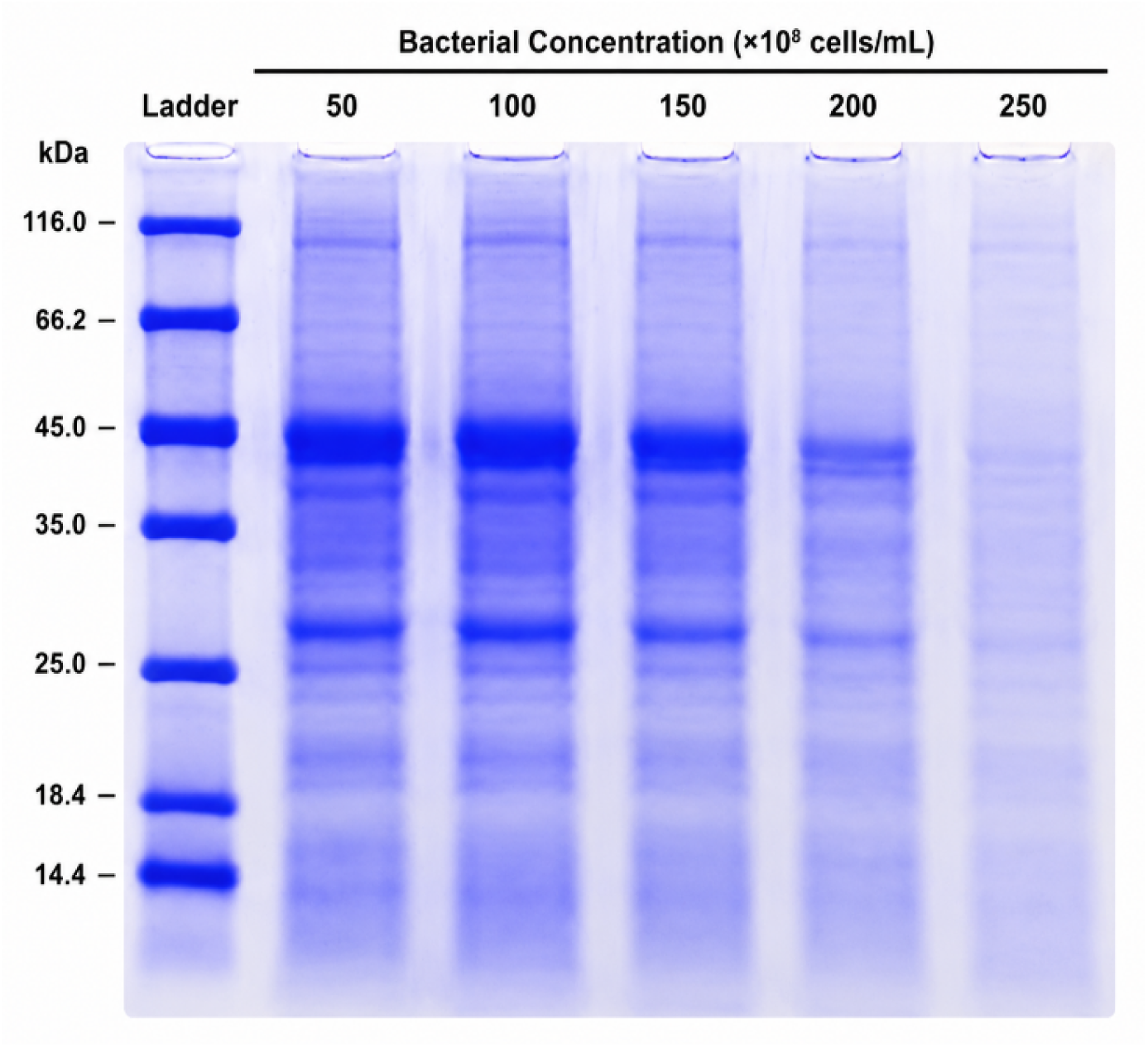
SDS-PAGE protein profile of *E. carotovora* after treatment with increasing concentrations of neem-mediated ZnO nanoparticles. Progressive changes in the bacterial protein profile are visible across treatment lanes.

#### 3.2.4 Antioxidant activity

The antioxidant activity of neem-mediated ZnO nanoparticles was evaluated using the DPPH radical scavenging assay (Fig. 7). A concentration-dependent increase in DPPH radical scavenging activity was observed for neem-mediated ZnO nanoparticles, with values increasing from 47.1% (T1) to 58.2% (T2), 71.0% (T3), and 89.4% (T4). The positive control (T5) showed a radical scavenging activity of 94.5%. Linear regression analysis demonstrated a strong concentration-response relationship, indicating a significant increase in radical scavenging activity with increasing nanoparticle concentration.

**Fig. 7.**
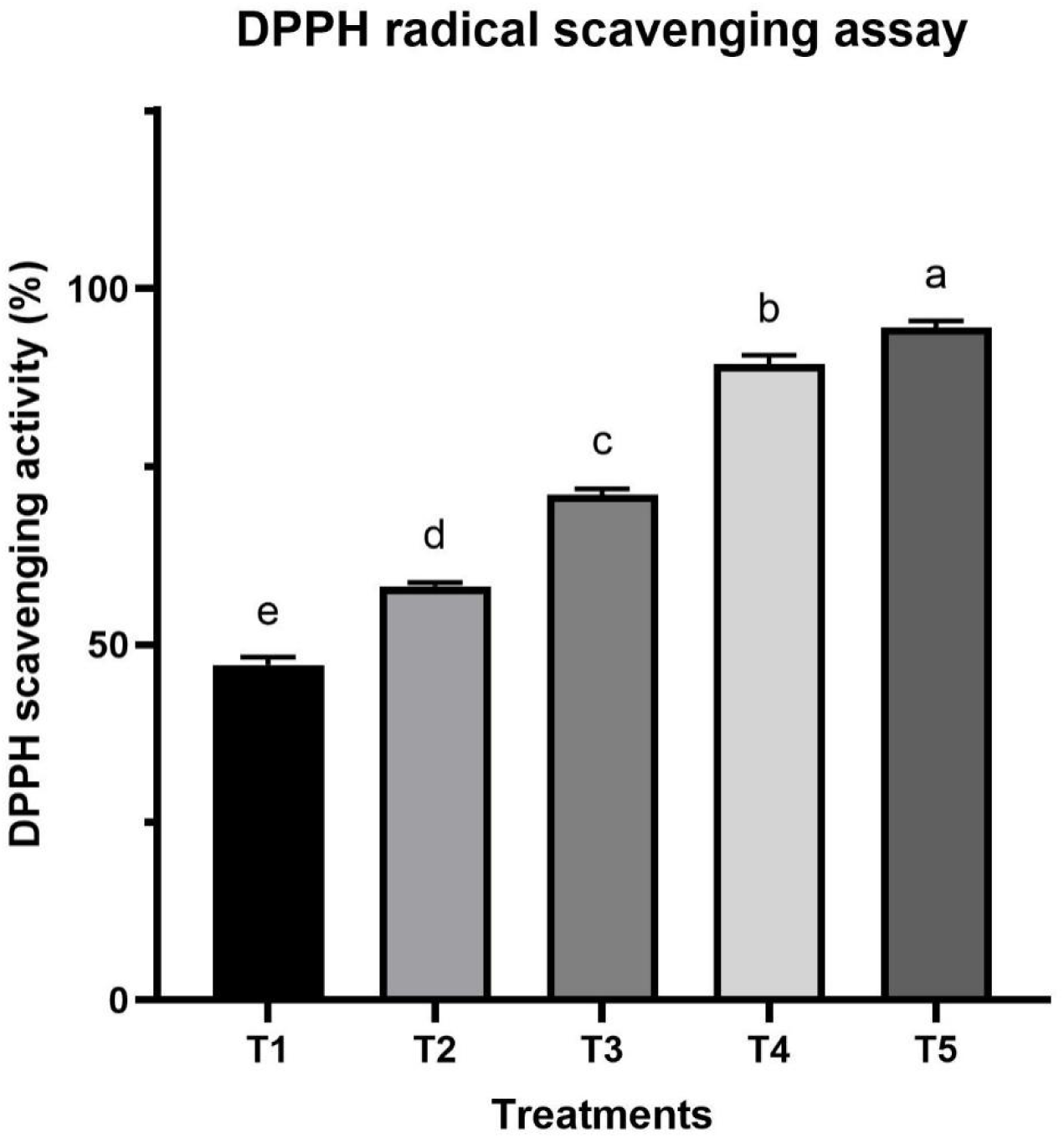
DPPH radical scavenging activity of neem-mediated ZnO nanoparticles at different concentrations. T1-T4 correspond to ZnO nanoparticle concentrations of 50, 100, 150, and 200 μg/mL, respectively, and T5 represents the positive control (gentamicin, 50 μg/well). Bars represent the mean DPPH radical scavenging activity (%) ± SD (n = 3). Different lowercase letters indicate statistically significant differences among treatments (P < 0.05, one-way ANOVA followed by Tukey’s HSD test).

#### 3.2.5 Cytotoxicity Assay

The cytotoxic effects of neem-mediated ZnO nanoparticles were evaluated using HepG2 cells by the MTT assay (Fig. 8). Cell viability decreased in a concentration-dependent manner from 70.7% at 50 μg/mL (T1) to 44.7%, 18.7%, and 3.3% at 100 (T2), 150 (T3), and 200 μg/mL (T4), respectively. Nonlinear regression analysis of the dose-response curve estimated an IC_50_ value of 124.8 μg/mL. The positive control (T5) was analyzed separately and was not included in the IC₅₀ determination. Statistical analysis showed significant differences among treatment groups.

**Fig. 8.**
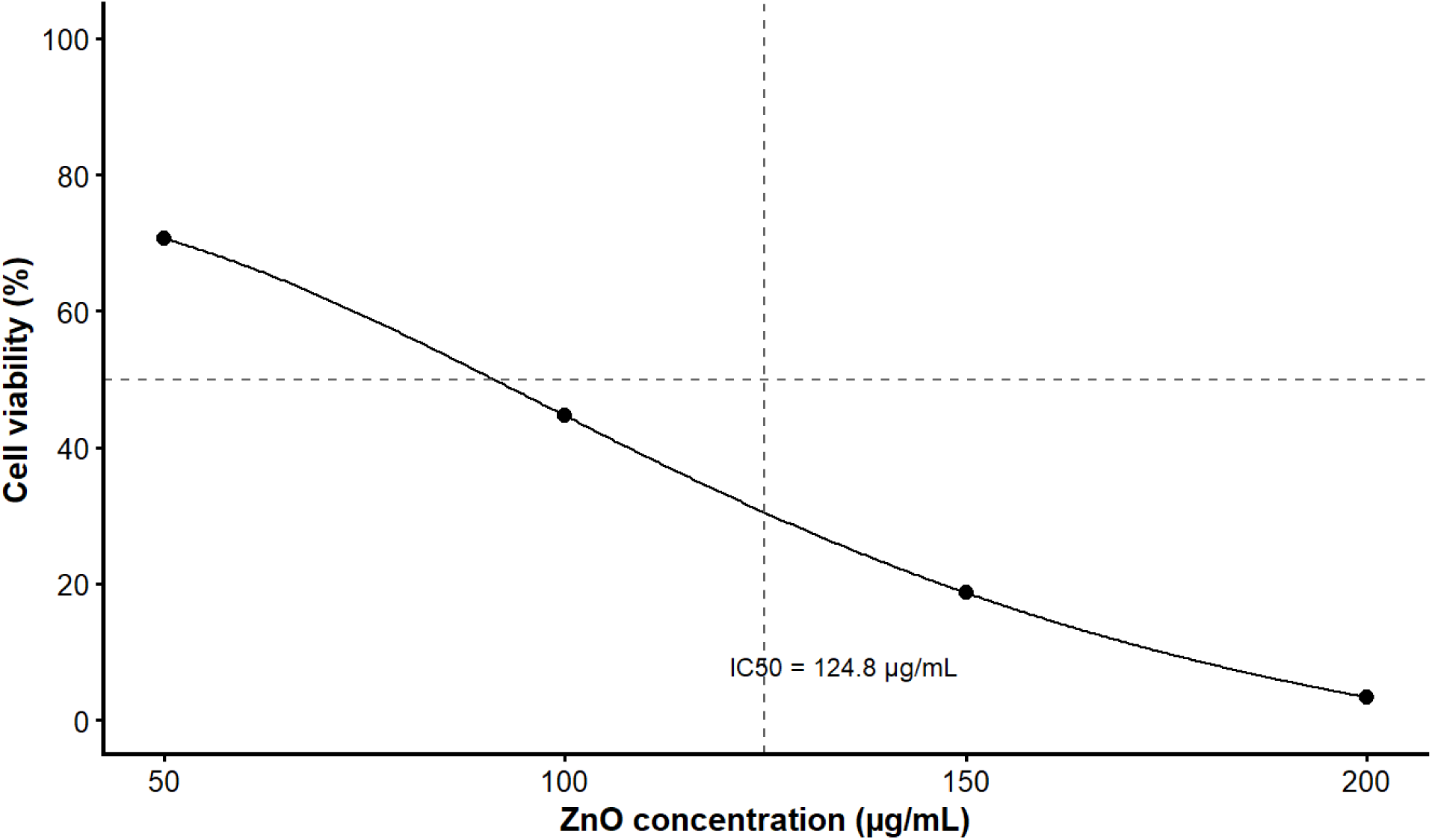
Cytotoxicity of neem-mediated ZnO nanoparticles toward HepG2 cells determined by the MTT assay. T1-T4 correspond to ZnO nanoparticle concentrations of 50, 100, 150, and 200 μg/mL, respectively, and T5 represents the positive control (gentamicin, 50 μg/well). Bars represent the mean cell viability (%) ± SD (n = 3). Different lowercase letters indicate statistically significant differences among treatments (P < 0.05, one-way ANOVA followed by Tukey’s HSD test).

## 4. Discussion

The successful synthesis of neem-mediated ZnO nanoparticles was confirmed through complementary optical, structural and chemical characterization. The characteristic UV–Visible absorption peak at 352 nm and the optical band gap of 3.07 eV are consistent with values commonly reported for green-synthesized ZnO nanoparticles. For example, Jaishi et al., (2024) reported an absorption maximum at 361 nm for *Moringa oleifera*-mediated ZnO nanoparticles, while Liga et al. (2024) observed a peak at 340 nm using isoflavone puerarin. Comparable absorption peaks have also been reported at 362 nm for flaxseed-mediated ZnO nanoparticles (Pehlivanoglu et al., 2023; Farooq U et al. 2022), 363 nm for pomegranate peel-mediated ZnO nanoparticles (Kapare et al., 2025), and 364 nm for *Wodyetia bifurcata*-mediated ZnO nanoparticles (Moalwi et al., 2024), confirming the successful formation of ZnO nanoparticles. Similarly, the optical band gap obtained in this study falls within the 3.07-3.34 eV range reported for plant-mediated ZnO nanoparticles (Kapare et al., 2025; Liga et al., 2024; López-López et al., 2021; Mutukwa et al., 2024; Raza et al., 2024). Minor variations among studies are expected because precursor concentration, reaction pH, phytochemical composition and crystallite size all influence the optical properties of ZnO nanoparticles. FTIR analysis demonstrated the presence of hydroxyl, carbonyl and C–O functional groups together with the characteristic Zn–O stretching vibration, indicating that neem-derived biomolecules participated in nanoparticle formation. Similar FTIR signatures have been reported for ZnO nanoparticles synthesized using *Azadirachta indica* and other medicinal plants, where phenolics and flavonoids function as natural reducing and capping agents and the ZnO peak observed under 550 cm^-1^ (Halder et al., 2025; Mohanasundaram & Saral A., 2025; Tsegahun & Aklilu, 2025). Although FTIR cannot identify individual phytochemicals, the observed functional groups support the presence of surface-associated biomolecules that may improve nanoparticle stability and influence subsequent biological interactions. XRD further confirmed the formation of phase-pure hexagonal wurtzite ZnO with an average crystallite size of 32 nm, which is comparable to the 28–35 nm reported by Halder et al. (2025) and the ∼30 nm crystallites described by Jaishi et al. (2024). GC–MS analysis identified 17 phytochemical constituents, demonstrating that a chemically diverse mixture of plant-derived compounds remained associated with the nanoparticle preparation and it is consistent with previous reports where GC–MS profiling of neem-mediated ZnO nanoparticles revealed 21 bioactive metabolites involved in nanoparticle stabilization and surface functionalization (El-Beltagi et al., 2024). Rather than attributing biological activity to individual metabolites identified by NIST library matching, which remains tentative, the collective phytochemical coating is more likely to contribute to nanoparticle stabilization and modify interactions with bacterial cells.

The favourable physicochemical characteristics of the synthesized ZnO nanoparticles translated into strong antibacterial activity against *E. carotovora*. The inhibition zone increased from 9.17 ± 0.60 mm at the lowest nanoparticle concentration to 27.20 ± 0.70 mm at the highest concentration, demonstrating a clear dose-dependent response. Previous studies have demonstrated the broad-spectrum antibacterial activity of green-synthesized ZnO nanoparticles against phytopathogenic bacteria. Cheema et al. (2022) reported a maximum inhibition zone of 25.7 mm against *Xanthomonas oryzae* pv. *oryzae* at 20 µg/mL ZnO nanoparticles, whereas neem-functionalized ZnO nanoparticles showed strong antibacterial activity against *Bacillus subtilis* and *Staphylococcus aureus* at 50 and 400 µg/mL, respectively (Halder et al., 2025). Similarly, Khan et al. (2021) demonstrated strong antibacterial activity against *Ralstonia solanacearum*, where ZnO nanoparticles produced a 22.3 mm inhibition zone at 18 µg/mL (Khan et al., 2021; Etminani F et al. 2023). The differences in inhibition zones between the present study and previous studies most likely arise from differences in plant extract chemistry, nanoparticle physicochemical properties, bacterial species and experimental methodology. Recent reviews have emphasized that the antibacterial performance of biogenic ZnO nanoparticles is governed not only by particle size but also by crystallinity, surface charge, phytochemical capping, morphology, synthesis conditions and nanoparticle dispersion (Okaiyeto et al., 2024). These studies provided evidence that the nanoparticles synthesized from different plant species frequently exhibit different antibacterial efficiencies even when their crystallite sizes are comparable.

The mechanism underlying bacterial inhibition was investigated through membrane integrity using Bradford and ninhydrin assays. ZnO nanoparticle treatment resulted in an approximately 4.8-fold increase in extracellular protein leakage and a 5.6-fold increase in amino acid leakage compared with untreated cells. These findings agree with previous reports showing that ZnO nanoparticles increase bacterial membrane permeability, resulting in leakage of intracellular biomolecules following nanoparticle exposure (Sirelkhatim et al., 2015). Recent mechanistic studies by Elabbasy et al. (2025) further demonstrated that green-synthesized ZnO nanoparticles disrupt bacterial membranes, alter cellular morphology and interfere with intracellular integrity. The leakage increased progressively during the first 12 h of exposure before reaching a plateau, indicating extensive membrane disruption after 12 hours the change was not statistically significant. Similarly, Gunalan et al. (2012) reported concentration-dependent protein leakage following green ZnO nanoparticle treatment, with extracellular protein levels reaching approximately 280 µg/mL in *S. aureus* and 250 µg/mL in *P. mirabilis* after 12 h. After 24 h of treatment at 8× MIC, protein leakage reached approximately 1.4 mg/mL in *S. aureus*, 1.2 mg/mL in *E. coli*, and 1.0 mg/mL in *Salmonella* spp., confirming that ZnO nanoparticles compromise bacterial membrane integrity (Mohd Yusof et al., 2021). The greater increase in amino acid leakage and protein leakage observed in the present study provides additional insight into the antibacterial process. The membrane permeabilization occurred progressively rather than through immediate cell lysis, an interpretation that is consistent with previous reports describing gradual membrane destabilization during ZnO nanoparticle exposure.

SDS–PAGE analysis further strengthened this interpretation by revealing progressive reductions in bacterial protein band intensity within the 29–66 kDa molecular weight range following nanoparticle treatment. Reduced intensity of several protein bands following ZnO nanoparticle treatment is consistent with Zanet et al. (2019), who observed marked changes in SDS–PAGE protein profiles, including decreased expression of proteins involved in amino acid metabolism and protein synthesis. Other than protein profiling, the antioxidant evaluation demonstrated similar concentration-dependent free radical scavenging activity, with DPPH inhibition increasing progressively from 47.1% to 75.0%. The previous studies also reported a concentration-dependent increase where Cheah et al. (2026) reported a maximum DPPH radical scavenging activity of 62.47% for pomegranate husk-mediated ZnO nanoparticles after 60 min of incubation, whereas Sangeetha & Hemamalini, (2025) observed a maximum DPPH scavenging activity of 89.0% for clove- and cinnamon-mediated ZnO nanoparticles. Similarly, Halfeld et al., (2026) reported that *Myrcia oblongata*-derived ZnO nanoparticles possessed more than twofold higher antioxidant capacity than commercial ZnO nanoparticles. These differences between different green synthesized nanoparticles are strongly influenced by the phytochemical composition of the plant extract, nanoparticle surface chemistry, extraction procedure and synthesis conditions. Neem leaves contain diverse flavonoids, phenolic compounds, limonoids, and tannins that participate in nanoparticle synthesis. In contrast, pomegranate husk contains abundant ellagitannins and anthocyanins, whereas clove and cinnamon are particularly rich in eugenol, cinnamaldehyde and related phenylpropanoids, all of which differ in antioxidant potential and surface interaction with ZnO nanoparticles.

MTT assay further demonstrated that the biological response of HepG2 cells was concentration dependent, with cell viability decreasing from 70% at the lowest nanoparticle concentration to 3% at the highest concentration with estimated IC_50_ value of 124.8 μg/mL. Sangeetha & Hemamalini, (2025) observed cytotoxicity against HepG2 cells, reporting an IC₅₀ value of 147.44 μg/mL, suggesting that plant-mediated ZnO nanoparticles maintain acceptable biocompatibility at lower concentrations while exhibiting greater biological activity at higher doses. While Yadav et al. (2025) and Nisha Jenifar et al. (2025) reported cytotoxicity of green-synthesized ZnO nanoparticles against HepG2 cells, with IC₅₀ values of 49.83 and 69.91 μg/mL, respectively, demonstrating progressive reductions in cell viability with increasing nanoparticle concentration. Each study exhibits specific dose dependent nanoparticle uptake efficiencies and metabolic responses with specific IC_50_ values which demonstrates that the plant metabolites contribute most against the biological activities of ZnO nanoparticles. This demonstrates that neem-mediated ZnO nanoparticles possess a balanced combination of antibacterial activity, antioxidant potential and cytocompatibility at lower concentrations. This integrated approach broadens the application of plant-mediated ZnO nanoparticles to sustainable agricultural disease management, highlighting their potential as eco-friendly alternatives for controlling pathogens. Nevertheless, the proposed antibacterial mechanism is based primarily on biochemical evidence, and future investigations combining intracellular ROS measurements, transcriptomic and proteomic analyses, together with greenhouse and field evaluations, will be essential to validate the molecular mechanisms and practical effectiveness of these nanoparticles under agricultural conditions.

## Conclusion

This study presents a sustainable strategy for the green synthesis of phytochemically functionalized ZnO nanoparticles using *Azadirachta indica* leaf extract and provides mechanistic evidence supporting their antibacterial activity against *E. carotovora*. The biosynthesized nanoparticles combined desirable physicochemical characteristics with potent concentration-dependent antibacterial activity, which was associated with membrane disruption, intracellular protein and amino acid leakage, and substantial alterations in bacterial protein profiles. Furthermore, their strong antioxidant activity and moderate cytocompatibility highlight the multifunctional nature of these nanomaterials. The integration of comprehensive physicochemical characterization with mechanistic antibacterial investigations provides new insights into the mode of action of plant-mediated ZnO nanoparticles and supports their potential as ecofriendly alternatives to conventional bactericides. Further validation under greenhouse and field conditions, together with studies on formulation stability and biosafety, will facilitate their use in sustainable crop protection strategies.

## Declarations

## Ethics approval and consent to participate

This study does not include any experiments with humans or other living animals, hence ethical approval and consent to participate is not applicable.

## Consent to publication

Not applicable.

## Data availability statement

This article contains experimental data generated or analyzed during the study. The corresponding author may provide additional information on reasonable request.

## Author’s Contributions

SM, TA and HR conceptualization, resources, analyses, writing, and methodology; SY, and SA, investigation, software, validation; review and editing, visualization, and supervision. All authors have carefully reviewed and consented to the final version of the manuscript.

## References

Al-darwesh, M. Y., Ibrahim, S. S., & Mohammed, M. A. (2024). A review on plant extract mediated green synthesis of zinc oxide nanoparticles and their biomedical applications. Results in Chemistry, 7, 101368. 10.1016/j.rechem.2024.101368

Bradford, M. M. (1976). A rapid and sensitive method for the quantitation of microgram quantities of protein utilizing the principle of protein-dye binding. Analytical Biochemistry, 72(1), 248–254. 10.1016/0003-2697(76)90527-3

Cheah, S.-Y., Phang, Y.-K., Lim, S. C.-Y., Koh, M.-X., Djearamane, S., Subramaniam, H., Lim, B.-H., Li, F., Aminuzzaman, M., Wong, L.-S., & Tey, L.-H. (2026). Green synthesis of ZnO nanoparticles using pomegranate husk extract: comparative evaluation of antioxidant, enzyme inhibition, and cytotoxic properties. Artificial Cells, Nanomedicine, and Biotechnology, 54(1), 120–136. 10.1080/21691401.2026.2668258

Cheema, A. I., Ahmed, T., Abbas, A., Noman, M., Zubair, M., & Shahid, M. (2022). Antimicrobial activity of the biologically synthesized zinc oxide nanoparticles against important rice pathogens. Physiology and Molecular Biology of Plants : An International Journal of Functional Plant Biology, 28(10), 1955–1967. 10.1007/s12298-022-01251-y

Dey, S., Mohanty, D. lochan, Divya, N., Bakshi, V., Mohanty, A., Rath, D., Das, S., Mondal, A., Roy, S., & Sabui, R. (2025). A critical review on zinc oxide nanoparticles: Synthesis, properties and biomedical applications. Intelligent Pharmacy, 3(1), 53–70. 10.1016/j.ipha.2024.08.004

El-beltagi, H. S., Ragab, M., Osman, A., & El-masry, R. A. (2024). Biosynthesis of zinc oxide nanoparticles via neem extract and their anticancer and antibacterial activities. 1–33. 10.7717/peerj.17588

El-Beltagi, H. S., Ragab, M., Osman, A., El-Masry, R. A., Alwutayd, K. M., Althagafi, H., Alqahtani, L. S., Alazragi, R. S., Alhajri, A. S., & El-Saber, M. M. (2024). Biosynthesis of zinc oxide nanoparticles via neem extract and their anticancer and antibacterial activities. PeerJ, 12, e17588. 10.7717/peerj.17588

El-Saadony, M. T., Fang, G., Yan, S., Alkafaas, S. S., El Nasharty, M. A., Khedr, S. A., Hussien, A. M., Ghosh, S., Dladla, M., Elkafas, S. S., Ibrahim, E. H., Salem, H. M., Mosa, W. F. A., Ahmed, A. E., Mohammed, D. M., Korma, S. A., El-Tarabily, M. K., Saad, A. M., El-Tarabily, K. A., & AbuQamar, S. F. (2024). Green Synthesis of Zinc Oxide Nanoparticles: Preparation, Characterization, and Biomedical Applications - A Review. International Journal of Nanomedicine, 19, 12889–12937. 10.2147/IJN.S487188

Elabbasy, M. T., El Bayomi, R. M., Abdelkarim, E. A., Hafez, A. E.-S. E., Othman, M. S., Ghoniem, M. E., Samak, M. A., Alshammari, M. H., Almarshadi, F. A., Elsamahy, T., & Hussein, M. A. (2025). Harnessing Stevia rebaudiana for Zinc Oxide Nanoparticle Green Synthesis: A Sustainable Solution to Combat Multidrug-Resistant Bacterial Pathogens. Nanomaterials (Basel, Switzerland), 15(5). 10.3390/nano15050369

Fei, H., Zhang, X., Fremah, O. G., Godana, E. A., Li, J., Xie, Y., Zhao, L., & Zhang, H. (2026). Exploring the mechanisms involved in Pectobacterium carotovorum subsp. brasiliense infecting postharvest tomato fruits. Postharvest Biology and Technology, 234, 114136. 10.1016/j.postharvbio.2025.114136

Gunalan, S., Sivaraj, R., & Rajendran, V. (2012). Green synthesized ZnO nanoparticles against bacterial and fungal pathogens. Progress in Natural Science: Materials International, 22(6), 693–700. 10.1016/j.pnsc.2012.11.015

Halder, A., Mohan, G. R., Matheshwaran, S., & Jha, S. K. (2025). Green synthesis of neem (Azadirachta indica) functionalized zinc oxide with enhanced antimicrobial properties. Next Materials, 8, 100725. 10.1016/j.nxmate.2025.100725

Halfeld, M., Rosset, J., Cinel, V. D. P., Aragão, C. B., Mariano, K. C. F., Nunes, R. S., Battistini, G. C., da Silva, R. A. G., Pinto, F. G. S., Seabra, A. B., & dos Reis, R. A. (2026). Green Synthesis of ZnO Nanoparticles Using Myrcia Oblongata DC: An Ecological Alternative for Biomedical Applications. Journal of Cluster Science, 37(2), 49. 10.1007/s10876-026-03000-7

Jaishi, D. R., Ojha, I., Bhattarai, G., Baraili, R., Pathak, I., Ojha, D. R., Shrestha, D. K., & Sharma, K. R. (2024). Plant-mediated synthesis of zinc oxide (ZnO) nanoparticles using Alnus nepalensis D. Don for biological applications. Heliyon, 10(20). 10.1016/j.heliyon.2024.e39255

Kapare, H. S., Bhosale, M., Karwa, P., Kulkarni, D., Bhole, R., & Labhade, S. (2025). Phyto-Assisted Synthesis and Investigation of Zinc Oxide Nanoparticles for Their Anti-Aging, Sun Protection and Antibacterial Activity. In Cosmetics (Vol. 12, Issue 6, p. 238). 10.3390/cosmetics12060238

Khalil, H., & Villota, R. (1988). Comparative Study on Injury and Recovery of Staphylococcus aureus using Microwaves and Conventional Heating. Journal of Food Protection, 51(3), 181–186. 10.4315/0362-028X-51.3.181

Khan, R. A. A., Tang, Y., Naz, I., Alam, S. S., Wang, W., Ahmad, M., Najeeb, S., Rao, C., Li, Y., Xie, B., & Li, Y. (2021). Management of Ralstonia solanacearum in Tomato Using ZnO Nanoparticles Synthesized Through Matricaria chamomilla. Plant Disease, 105(10), 3224– 3230. 10.1094/PDIS-08-20-1763-RE

Lebaka, V. R., Ravi, P., Reddy, M. C., Thummala, C., & Mandal, T. K. (2025). Zinc Oxide Nanoparticles in Modern Science and Technology: Multifunctional Roles in Healthcare, Environmental Remediation, and Industry. Nanomaterials (Basel, Switzerland), 15(10). 10.3390/nano15100754

Liga, S., Vodă, R., Lupa, L., Paul, C., Nemeş, N. S., Muntean, D., Avram, Ștefana, Gherban, M., & Péter, F. (2024). Green Synthesis of Zinc Oxide Nanoparticles Using Puerarin: Characterization, Antimicrobial Potential, Angiogenesis, and In Ovo Safety Profile Assessment. Pharmaceutics, 16(11). 10.3390/pharmaceutics16111464

López-López, J., Tejeda-Ochoa, A., López-Beltrán, A., Herrera-Ramírez, J., & Méndez-Herrera, P. (2021). Sunlight Photocatalytic Performance of ZnO Nanoparticles Synthesized by Green Chemistry Using Different Botanical Extracts and Zinc Acetate as a Precursor. Molecules (Basel, Switzerland), 27(1). 10.3390/molecules27010006

Moalwi, A., Kamat, K., Muddapur, U. M., Aldoah, B., AlWadai, H. H., Alamri, A. M., Alrashid, F. F., Alsareii, S. A., Mahnashi, M. H., Shaikh, I. A., Khan, A. A., & More, S. S. (2024). Green synthesis of zinc oxide nanoparticles from Wodyetia bifurcata fruit peel extract: multifaceted potential in wound healing, antimicrobial, antioxidant, and anticancer applications. Frontiers in Pharmacology, Volume 15-2024. https://www.frontiersin.org/journals/pharmacology/articles/10.3389/fphar.2024.1435222

Mohammed, A. M., Mohammed, M., Oleiwi, J. K., Ihmedee, F. H., Adam, T., Betar, B. O., & Gopinath, S. C. B. (2025). Comprehensive review on zinc oxide nanoparticle production and the associated antibacterial mechanisms and therapeutic potential. Nano Trends, 11, 100145. 10.1016/j.nwnano.2025.100145

Mohanasundaram, P., & Saral A., M. (2025). Binding properties and biological applications of green synthesized ZnO nanoparticles from neem flower. Scientific Reports, 15(1), 17727. 10.1038/s41598-025-02157-x

Mohd Yusof, H., Abdul Rahman, N., Mohamad, R., Hasanah Zaidan, U., & Samsudin, A. A. (2021). Antibacterial Potential of Biosynthesized Zinc Oxide Nanoparticles against Poultry-Associated Foodborne Pathogens: An In Vitro Study. Animals : An Open Access Journal from MDPI, 11(7). 10.3390/ani11072093

Mosmann, T. (1983). Rapid colorimetric assay for cellular growth and survival: Application to proliferation and cytotoxicity assays. Journal of Immunological Methods, 65(1), 55–63. 10.1016/0022-1759(83)90303-4

Mutukwa, D., Taziwa, R. T., Tichapondwa, S. M., & Khotseng, L. (2024). Optimisation, Synthesis, and Characterisation of ZnO Nanoparticles Using Leonotis ocymifolia (L. ocymifolia) Leaf Extracts for Antibacterial and Photodegradation Applications. International Journal of Molecular Sciences, 25(21). 10.3390/ijms252111621

Nan, J., Chu, Y., Guo, R., & Chen, P. (2024). Research on the antibacterial properties of nanoscale zinc oxide particles comprehensive review. Frontiers in Materials, *Volume* 11-2024. https://www.frontiersin.org/journals/materials/articles/10.3389/fmats.2024.1449614

Nisha Jenifar, A., Anilkumar, P., & Preetha, S. (2025). Eco-friendly fabrication of ZnO nanocomposites using Lepidium didymium and polymer-assisted CTAB/PEG: A multifunctional approach for enhanced biomedical applications. Journal of Molecular Liquids, 417, 126550. 10.1016/j.molliq.2024.126550

Okaiyeto, K., Gigliobianco, M. R., & Martino, P. Di. (2024). Biogenic Zinc Oxide Nanoparticles as a Promising Antibacterial Agent : Synthesis and Characterization.

Pachaiappan, R., Rajendran, S., Ramalingam, G., Vo, D.-V. N., Priya, P. M., & Soto-Moscoso, M. (2021). Green Synthesis of Zinc Oxide Nanoparticles by Justicia adhatoda Leaves and Their Antimicrobial Activity. Chemical Engineering & Technology, 44(3), 551–558. 10.1002/ceat.202000470

Pehlivanoglu, S., Acar, C. A., & Donmez, S. (2023). Characterization of green synthesized flaxseed zinc oxide nanoparticles and their cytotoxic, apoptotic and antimigratory activities on aggressive human cancer cells. Inorganic and Nano-Metal Chemistry, 53(9), 1022–1031. 10.1080/24701556.2021.1980034

Perfileva, A. I., Strekalovskaya, E. I., Klushina, N. V, Gorbenko, I. V, & Krutovsky, K. V. (2025). The Causative Agent of Soft Rot in Plants, the Phytopathogenic Bacterium Pectobacterium carotovorum subsp. carotovorum: A Brief Description and an Overview of Methods to Control It. In Agronomy (Vol. 15, Issue 7, p. 1578). 10.3390/agronomy15071578

Rahman, F., Majed Patwary, M. A., Bakar Siddique, M. A., Bashar, M. S., Haque, M. A., Akter, B., Rashid, R., Haque, M. A., & Royhan Uddin, A. K. M. (2022). Green synthesis of zinc oxide nanoparticles using Cocos nucifera leaf extract: characterization, antimicrobial, antioxidant and photocatalytic activity. Royal Society Open Science, 9(11), 220858. 10.1098/rsos.220858

Rani, N., Sagar, N. A., Chauhan, A., & Mondal, A. (2025). Green synthesis of ZnO nanoparticles: Characterization and emerging applications in sustainable agriculture. Industrial Crops and Products, 233, 121393. 10.1016/j.indcrop.2025.121393

Raza, A., Malan, P., Ahmad, I., Khan, A., Haris, M., Zahid, Z., Jameel, M., Ahmad, A., Seth, C. S., Asseri, T. A. Y., Hashem, M., & Ahmad, F. (2024). Polyalthia longifolia-mediated green synthesis of zinc oxide nanoparticles: characterization, photocatalytic and antifungal activities. RSC Advances, 14(25), 17535–17546. 10.1039/d4ra01035c

Sangeetha, V., & Hemamalini, A. J. (2025). Sustainable Green Synthesis of Zinc Oxide Nanoparticles with Clove and Cinnamon : A Study on Antioxidant and Cytotoxic Properties. 19(4), 257–262.

Sarkar, S., Singh, R. P., & Bhattacharya, G. (2021). Exploring the role of Azadirachta indica (neem) and its active compounds in the regulation of biological pathways: an update on molecular approach. 3 Biotech, 11(4), 178. 10.1007/s13205-021-02745-4

Sharma, Y., Anand, V., Kumar, R., Kumar, A., & Heera, P. (2025). Green synthesized ZnO nanoparticles using Jatropha curcas latex for antibacterial applications. Next Materials, 8, 100869. 10.1016/j.nxmate.2025.100869

Sirelkhatim, A., Mahmud, S., Seeni, A., Kaus, N. H. M., Ann, L. C., Bakhori, S. K. M., Hasan, H., & Mohamad, D. (2015). Review on Zinc Oxide Nanoparticles: Antibacterial Activity and Toxicity Mechanism. Nano-Micro Letters, 7(3), 219–242. 10.1007/s40820-015-0040-x

Swain, M., Mishra, D., & Sahoo, G. (2025). A review on green synthesis of ZnO nanoparticles. Discover Applied Sciences, 7(9), 997. 10.1007/s42452-025-06957-8

Takcı, D. K., Ozdenefe, M. S., Huner, T., & Takcı, H. A. M. (2025). Plant-mediated green route to the synthesis of zinc oxide nanoparticles: in vitro antibacterial potential. Journal of the Australian Ceramic Society, 61(1), 31–39. 10.1007/s41779-024-01064-0

Tsegahun, E., & Aklilu, M. (2025). Neem (Azadirachta indica) leaf extract mediated synthesis of zinc oxide nanoparticles (ZnO NPs) and their antibacterial activity. Discover Nano, 20(1), 145. 10.1186/s11671-025-04260-4

Villagrán, Z., Anaya-Esparza, L. M., Velázquez-Carriles, C. A., Silva-Jara, J. M., Ruvalcaba-Gómez, J. M., Aurora-Vigo, E. F., Rodríguez-Lafitte, E., Rodríguez-Barajas, N., Balderas-León, I., & Martínez-Esquivias, F. (2024). Plant-Based Extracts as Reducing, Capping, and Stabilizing Agents for the Green Synthesis of Inorganic Nanoparticles. In Resources (Vol. 13, Issue 6, p. 70). 10.3390/resources13060070

Wahab, R., Siddiqui, M. A., Saquib, Q., Dwivedi, S., Ahmad, J., Musarrat, J., Al-Khedhairy, A. A., & Shin, H.-S. (2014). ZnO nanoparticles induced oxidative stress and apoptosis in HepG2 and MCF-7 cancer cells and their antibacterial activity. Colloids and Surfaces B: Biointerfaces, 117, 267–276. 10.1016/j.colsurfb.2014.02.038

Yadav, A., Kumar, H., Kumar, P., Rani, G., & Maken, S. (2025). Syzygium cumini leaf extract mediated green synthesis of ZnO nanoparticles: A sustained release for anticancer, antimicrobial, antioxidant, and anti-corrosive applications. Journal of Molecular Structure, 1325, 141017. 10.1016/j.molstruc.2024.141017

Yagoub, A. E. A., Al-Shammari, G. M., Al-Harbi, L. N., Subash-Babu, P., Elsayim, R., Mohammed, M. A., Yahya, M. A., & Fattiny, S. Z. A. (2022). Antimicrobial Properties of Zinc Oxide Nanoparticles Synthesized from Lavandula pubescens Shoot Methanol Extract. In Applied Sciences (Vol. 12, Issue 22, p. 11613). 10.3390/app122211613

Zanet, V., Vidic, J., Auger, S., Vizzini, P., Lippe, G., Iacumin, L., Comi, G., & Manzano, M. (2019). Activity evaluation of pure and doped zinc oxide nanoparticles against bacterial pathogens and Saccharomyces cerevisiae. Journal of Applied Microbiology, 127(5), 1391– 1402. 10.1111/jam.14407

Etminani F, Etminani A, X Hasson SO, Kareem Judi H, Akter S, Saki M. (2023). In silico study of inhibition effects of phytocompounds from four medicinal plants against the Staphylococcus aureus β-lactamase. Informatics in Medicine Unlocked. 37, 2023, 101186. 10.1016/j.imu.2023.101186

Farooq U; Akter S; Qureshi AQ; Hayaa M. A; Farzana M; Shahab M et al. Arbutin Stabilized Silver Nanoparticles: Synthesis, Characterization, and Its Catalytic Activity against Different Organic Dyes. Catalysts 2022, Volume 12, Issue 12, 1602

